# Bayesian marker-based principal component ridge regression – a flexible multipurpose framework for quantitative genetics in wild study systems

**DOI:** 10.1101/2024.06.01.596874

**Authors:** Janne C. H. Aspheim, Kenneth Aase, Geir H. Bolstad, Henrik Jensen, Stefanie Muff

## Abstract

As larger genomic data sets become available for wild study populations, the need for flexible and efficient methods to estimate and predict quantitative genetic parameters, such as the adaptive potential and measures for genetic change, increases. Animal breeders have produced a wealth of methods, but wild study systems often face challenges due to larger effective population sizes, environmental heterogeneity and higher spatio-temporal variation. Here we adapt methods previously used for genomic prediction in animal breeding to the needs of wild study systems. The core idea is to approximate the breeding values as a linear combination of principal components (PCs), where the PC effects are shrunk with Bayesian ridge regression. Thanks to efficient implementation in a Bayesian framework using integrated nested Laplace approximations (INLA), it is possible to handle models that include several fixed and random effects in addition to the breeding values. Applications to a Norwegian house sparrow meta-population, as well as simulations, show that this method efficiently estimates the additive genetic variance and accurately predicts the breeding values. A major benefit of this modeling framework is computational efficiency at large sample sizes. The method therefore suits both current and future needs to analyze genomic data from wild study systems.

## 1 Introduction

When aiming to understand how wild populations respond to environmental change, or whether the population is able to respond at all, quantitative genetic methods can provide crucial insight into the underlying adaptive evolutionary potential (Falconer and Mackay, 1996; Lynch and Walsh, 1998; Charmantier et al., 2014). For wild systems, quantitative genetics serves two key tasks. First, the estimation of the *additive genetic variance* (*V_A_*) within a population, which corresponds to the part of the variance of a phenotype in a population that is explained by the additive effects of genes (Lynch and Walsh, 1998). Second, the within-sample estimation and out-of-sample prediction of *breeding values* (the sum of additive effects of an individual’s genes), known as *genomic prediction* (Meuwissen et al., 2001; McGaugh et al., 2021). The *V_A_* for a trait of interest reflects the heritable genetic variation in a population, and as such it is a fundamental concept both in evolutionary biology (Walsh and Lynch, 2018; Hansen et al., 2023) and biodiversity conservation (Kardos et al., 2021). Estimates of *V_A_* in fitness or fitness-related traits enable us to predict evolution by natural selection (Lande, 1979; Lande and Arnold, 1983; Bonnet et al., 2022). Larger values of *V_A_* are associated with a higher potential to adapt to changing or new environments, thus estimates of *V_A_* can be used as an indicator for the viability and robustness of a population exposed to a changing environment (Carlson et al., 2014; Haddad et al., 2015). Genomic prediction methods, on the other hand, estimate and predict individual-specific breeding values, and thus enable us to monitor evolutionary change. This is crucial, for example for conservation efforts when individuals have to be selected for captive breeding or translocation, but also to quantify the rate and direction of micro-evolutionary changes over time or across space (Jensen et al., 2014; Hunter et al., 2022).

For both *V_A_* estimation and genomic prediction, relatedness information between any pairs of individuals is traditionally the key information that underlies the statistical models (Henderson, 1984; Kruuk, 2004). Both tasks have previously been tackled via the use of long-term pedigree data (*e.g.*, Charmantier et al., 2014; Kvalnes et al., 2017; Bonnet et al., 2019; Reid et al., 2021), but as the costs of generating single nucleotide polymorphsim (SNP) or other DNA variant data are decreasing, genomic data is generated at higher rates also for wild populations. When individual genomic data is available, evolutionary ecologists no longer need field data across many generations to reconstruct pedigrees to estimate additive genetic variances or breeding values. However, the successful use of genomic data crucially depends on suitable statistical methodology for wild systems. While the fields of plant and animal breeding and human genomics have brought up a wealth of data sets and analytical tools adjusted to their particular needs (*e.g.*, Meuwissen et al., 2001; García-Ruiz et al., 2016; Khera et al., 2018), the actual motivation, terminology and methods differ between the fields, and similarities and differences often remain under-recognized (Wray et al., 2019; McGaugh et al., 2021). In wild systems, particular methodological challenges occur because, in contrast to breeding systems, wild populations tend to be smaller and at the same time more complex due to uncontrollable environmental heterogeneity and demographic variation, as well as ongoing evolutionary processes such as selection, drift, or migration. Despite some promising applications based on genomic data from wild study systems (Gienapp et al., 2019; Stocks et al., 2019; Ashraf et al., 2022; Hunter et al., 2022; Aase et al., 2022), the existing statistical methods have not yet been broadly adapted to the peculiarities of wild systems.

In the case when relatedness information is derived from pedigrees of wild systems, models were originally fitted via restricted maximum likelihood (REML) or its Bayesian variants (*e.g.*, Steinsland and Jensen, 2010; Hadfield, 2010) from a model denoted as the *animal model* to derive estimates of *V_A_* and best linear unbiased predictions (BLUPS) of the breeding values (Henderson, 1984; Wilson et al., 2010). However, when the entries in the relatively sparse matrix encoding for *expected* relatedness from the pedigree are replaced by estimates of *realized* relatedness among individuals, the resulting genomic relatedness matrix (GRM) in the *genomic genomic animal model* is very dense (VanRaden, 2008; Yang et al., 2011; Speed and Balding, 2015). Furthermore, the GRM is usually no longer positive definite, which prohibits its inversion. Even though approximate remedies for these problems exist (VanRaden, 2008; Zaitlen et al., 2013), statistical model fitting procedures quickly become inefficient with growing numbers of genotyped individuals.

With the explicit information from the SNPs, an alternative is to formulate a regression model with explicit additive marker effects, where the breeding value is the sum of all the SNP effects (Meuwissen et al., 2001). Marker-based regression is equivalent to the genomic animal model under appropriate standardization of the SNP genotypes (Habier et al., 2007; VanRaden, 2008; Goddard, 2009), and it has become popular in the past two decades, especially for genomic selection in animal and plant breeding, but more recently also for wild systems (*e.g.*, Meuwissen et al., 2016; Hickey et al., 2017; Ashraf et al., 2022; Hunter et al., 2022). However, even though a relatively small number of SNPs has shown to be sufficient for accurate estimation and prediction of parameters of interest (Bérénos et al., 2014; Kriaridou et al., 2020), the number of markers *m* usually greatly exceeds the often modest number of individuals *N* (“*N ≪ m*” problem), especially in wild systems. Simple regression is therefore not suitable to estimate the SNP-specific effects, and regularization techniques like BayesA, BayesB, BayesR, or the Bayesian LASSO (Meuwissen et al., 2001; Park and Casella, 2008; Habier et al., 2011; Erbe et al., 2012; Gianola, 2013; Moser et al., 2015) are needed. A major advantage of marker-based regression is that the computational complexity grows linearly with the number of individuals (even though as bas as cubic in the number of markers), while the size of the GRM in a genomic animal model – and thus the effort for matrix inversion – grows with its square (but linearly in the number of markers). On the other hand, existing methodology to fit explicit marker models was mainly developed for the large and highly inbred systems in homogeneous, controlled environments of animal and plant breeding contexts, and not for the complex wild study systems in non-constant environments often exposed to fluctuating selection and stochastic genetic processes (Bell, 2010). In fact, when applying existing methods such as BayesB or BayesR, researchers tend to either take simplifying assumptions, or implement a two-step strategy where environmental effects are accounted for by fitting models with the non-genetic effects first, and then using a suitable type of residuals in the actual genomic prediction analysis (*e.g.*, Ashraf et al., 2022; Hunter et al., 2022; Vahedi et al., 2023). However, the consequences of these simplifications have not been critically investigated, and it is for example unclear how they affect the estimators for *V_A_*, or the accuracy of the predicted breeding values.

Another way to handle the *N ≪ m* problem is via dimension reduction techniques. A particularly promising approach seems to be a combination of principal component (PC) and ridge-regression, with PCs derived from singular value decomposition (SVD) of the SNP matrix, where it can be assumed that the PC-effects are independent and normally distributed with the same variance (Ødegård et al., 2018). Selecting an appropriate prior distribution for the PC-effects is thus much easier than assigning an appropriate prior directly to the SNP effects, as needs to be done in the common regularization approaches (Gianola, 2013). The so-called principal component ridge regression (PCRR) method accurately and efficiently predicted breeding values in a large, homogeneous and highly inbred cattle population (Ødegård et al., 2018).

However, while a relatively small number of PCs is expected to explain large parts of the genetic variation in breeding systems that have small effective population size (*N_e_*) due to high degrees of relatedness (Hall, 2016), it is less clear how much wild systems benefit from the same approach. The generally larger *N_e_* (Palstra and Fraser, 2012), potential sub-structure (Wolak and Reid, 2017; Aase et al., 2022) and the fact that complex traits probably are affected by more genes than originally thought (Bulik-Sullivan et al., 2015; Goddard et al., 2016) indicates that a higher number of PCs is needed to obtain accurate models for wild systems. Moreover, previous applications of PC regression in the context of animal breeding did not account for additional fixed and random variables (Dadousis et al., 2014; Ødegård et al., 2018), which is a major limitation for the application in wild systems, where heterogeneous environmental conditions usually must be accounted for in order to obtain valid inference and good predictions.

Here we propose a flexible and general Bayesian version of the PCRR approach, denoted as BPCRR, which is able to handle in one single model all fixed and random effects that are necessary to account for the complexity in wild systems. An attractive feature of the method is that it can be used to address both key tasks of wild systems, namely for estimating *V_A_* and to do genomic prediction. To investigate the potential of BPCRR for the application in wild systems, we compare the method to the genomic animal model and BayesR in terms of computation times, unbiasedness of *V_A_* estimation and accuracy of genomic prediction, both for simulated data and for three complex traits (body mass, tarsus length and wing length) in a data set from an insular meta-population of house sparrows (*Passer domesticus*) in Northern Norway. The genomic animal model and BPCRR were treated in a Bayesian framework using integrated nested Laplace approximations (INLA) (Rue et al., 2009) via the R-interface `R-INLA` (Martins et al., 2013; Rue et al., 2017). BayesR models were fitted by taking advantage of the recent hibayes R package (Yin et al., 2022), a sampling-based Bayesian approach that now offers the possibility to fit models including any number of fixed and random effects and thus does not require a two-step procedure like previous implementations. To facilitate the use and implementation of the BPCRR approach, as well as the comparison to BayesR, we provide a coded example for a simulated case.

## 2 Methods

### 2.1 Statistical modeling background

#### 2.1.1 The animal model and the (G)BLUP

Variation in phenotypes of wild populations has in the past two decades traditionally been additively decomposed using the *animal model* (Henderson, 1976; Kruuk, 2004). This statistical modeling framework takes into account the relatedness between the studied individuals and decomposes the phenotype *y_i_* of an individual *i* in its most simple form into a genetic and an environmental component as

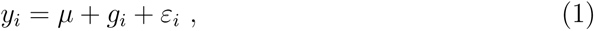

where *µ* is the population mean, *g_i_* is the additive genetic merit (breeding value) for individual *i*, and 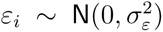 is the independent environmental effect. The vector of breeding values ***g****^⊤^* = (*g*_1_*, …, g_N_*) for *N* individuals is assumed to follow a multivariate normal distribution 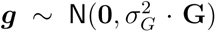, where **G** is the relatedness matrix and *σ_G_*^2^ the additive genetic variance, often equivalently denoted as *V_A_*. The relatedness matrix is either derived from pedigrees, leading to the pedigree-based animal model and predictions via the BLUP (Henderson, 1984), or, more recently, from genomic information, usually from individual SNP genotype data (VanRaden, 2008). In the latter case, the assumption is that marker effects in a large population are uncorrelated (VanRaden et al., 2009), and the **G** corresponds to the GRM. The predictions made by this genomic version of the animal model are commonly denoted as genomic BLUPS, or GBLUPs.

The animal model (1) can be extended in various ways in order to capture (permanent) environmental or individual-specific effects, like sex or age (Kruuk and Hadfield, 2007; Wilson et al., 2010). However, estimation tends to become challenging when additional environmental variation needs to be accounted for, namely when additional random effects need to be included, especially in the case where we also use the very dense GRM rather than the sparse pedigree-based relatedness matrix.

#### 2.1.2 Marker-based regression

In animal/plant breeding and human genomics, a popular alternative to (both the pedigree-based and genomic version of) the animal model (1) and its extensions is to formulate a regression model where the phenotype of interest explicitly depends on all the markers, and the breeding value *g_i_* is replaced by a sum over the effects of all genome-wide markers (*e.g.*, Meuwissen et al., 2001; Heffner et al., 2009; De Los Campos et al., 2010). This idea is based on the observation that quantitative traits in animals, plants and humans, such as body size, crop yield or disease status, are usually the result of many small contributions from loci across the genome, as well as fixed and environmental factors (Wood et al., 2014; Goddard et al., 2016; Walsh and Lynch, 2018). For a continuous trait, the *marker-based* linear model assumes that *m* SNPs for *N* individuals are available, and that (in the absence of repeated measurements) the *N ×* 1 vector of phenotypes ***y*** can be decomposed as a linear combination of contributions from all genomic markers summing up to the breeding value, plus any number of fixed and random effects as

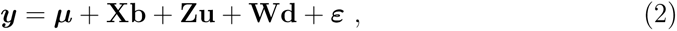

where ***µ*** is the vector for the overall mean, **X** is the matrix containing fixed-effect variables like sex or age with corresponding regression parameter vector **b**, **Z** is an *N × m* matrix containing the mean-centered marker codes (where the uncentered values typically are 0, 1 or 2 for the *AA*, *AB* and *BB* genotypes, respectively), and 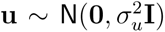 is a random effect for the allele substitution effects of each of the *m* markers, where **I** defines the identity matrix of appropriate dimension. Moreover, **W** is a design matrix of appropriate dimension for the random environmental effects **d**, and 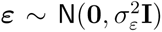 is the vector of normally distributed residuals. The vector of breeding values is then given as ***g*** = **Zu**. In the presence of repeated measurements for individuals, the dimensions in the components of model (2) are adjusted accordingly, and an individual-level *i.i.d.* random effect is added. Equivalence between the genomic animal model and SNP-based regression given by equation (2) holds when using the same SNPs, and assuming that effect sizes **u** all stem from the same distribution (Habier et al., 2007; VanRaden, 2008). In contrast to the genomic animal model, however, the main problem with model (2) is that we usually have more markers than individuals, that is, *N ≪ m*, thus a multiple linear regression approach cannot estimate the effect sizes of each SNP (*i.e.*, the elements in **u**). Standard least squares estimation is thus not feasible for model (2) and alternative approaches are thus needed.

#### 2.1.3 BayesR

A popular approach to handle the *N ≪ m* problem in marker-based regression is via Bayesian methods, where the idea is to assigning priors that reflect different hypotheses on the genetic architecture of a trait (Meuwissen et al., 2001; Habier et al., 2011; Erbe et al., 2012; Gianola, 2013; Moser et al., 2015). Imposing shrinkage through those priors allow us to obtain meaningful posterior distributions, even though the models are not likelihood identified. The members of the “Bayesian alphabet” have repeatedly been benchmarked against each other and against the genomic animal model and the (G)BLUP, in particular in terms of accuracy for genomic prediction (*e.g.*, Bolormaa et al., 2013; Duhnen et al., 2017; Mollandin et al., 2021; Ashraf et al., 2022; Meher et al., 2022). While the various methods have different strengths and weaknesses (Gianola, 2013; Meher et al., 2022), the BayesR approach suggested by Erbe et al. (2012) has consistently shown a competitive performance in terms of genomic prediction accuracy, while at the same time being relatively insensitive to the actual genetic architecture of the trait (*e.g.*, Kemper et al., 2015; Mollandin et al., 2021; Ashraf et al., 2022; Vahedi et al., 2023). The rationale behind BayesR is that most markers have no or a very small effect, and that the remaining markers affect the trait to different degrees, which is consistent with what is observed in practice (*e.g.*, Goddard et al., 2016; Yengo et al., 2022). The default BayesR prior on the effect sizes is a Gaussian mixture with zero mean and variances 0, 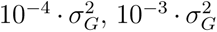 and 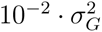, with Dirichlet priors on the probability vector ***π*** = (*π*_1_*, π*_2_*, π*_3_*, π*_4_) that weights the four components in the Gaussian mixture, and scaled inverse-chi squared distributions on *σ_G_*^2^ and *σ_ε_*^2^.

A challenge with BayesR is that its original implementation can neither handle repeated measurements, nor random effects other than the genetic value (Moser et al., 2015). The common work-around to fit models like (2) is to first pre-adjust the phenotypes for all fixed and random effects using a linear mixed model without a genetic component, and then use the individual-specific effect (in case of repeated measurements) or the residuals as the new response (Ashraf et al., 2022; Hunter et al., 2022; Vahedi et al., 2023). Even though prediction accuracy is usually quite high in those cases, a two-step approach is not very elegant, and it will break down if non-additive genetic effects or genotype-by-environment (*G × E*) interactions should be included. In addition, the procedure tends to significantly underestimate *σ_G_*^2^ (see Appendix A). Since we here are interested in estimating *σ_G_*^2^ and obtaining good predictions in the same framework, we rely on a recent implementation of BayesR in the R package hibayes (Yin et al., 2022), which can directly handle the full model (2). So far, the package has mostly been used for GWAS in animal- and plant breeding (Yang et al., 2021; Ding et al., 2022; Alboali et al., 2023) and in conservation biology (Guhlin et al., 2023), while only a few have tested it for genomic prediction (*e.g.*, Meher et al., 2023; Vahedi et al., 2023), and usually for relatively limited numbers of SNPs (*<* 18000 and *<* 47000 respectively). Here we will test hibayes the first time on a wild study system with *>* 180*^′^*000 SNPs.

### 2.2 The principal component ridge regression (BPCRR) approach

#### 2.2.1 Dimension reduction via singular value decomposition (SVD)

Another, so far under-explored avenue to tame the number of variables in marker-based regression in wild systems is via dimension reduction techniques (Ødegård et al., 2018; Hosseini-Vardanjani et al., 2018). In the latter case, the SNP matrix **Z** is decomposed via singular-value decomposition (SVD) **Z** = **USV***^⊤^*, where **U** has dimension *m × m*, **S** is a diagonal *m × N* matrix with singular values on the diagonal, and **V** is a *N × N* matrix of eigenvectors. The first *k* PCs of **Z** can then be extracted by multiplying the SNP matrix **Z** by the first *k* columns of **V** to obtain a matrix of reduced dimension **Z**★ = **ZV**[, 1: *k*], and reformulate model (2) as

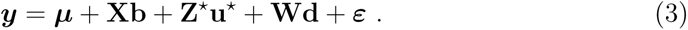

Instead of estimating *m* SNP effects, the problem is reduced to estimating the *k < m* PC-effects in 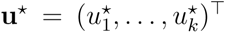, which is a computationally more manageable problem. Moreover, the orthogonality of the PCs imposes uncorrelatedness in the PC-effects, which is at the same time consistent with the assumption of uncorrelated marker effects in model (2) – an assumption that usually is violated for the SNP effects (Macciotta et al., 2010), and gets increasingly more violated when the number of SNPs increases.

Given that substantial linkage disequilibrium (LD) is expected in dense genomic data, and because related animals share alleles at a high proportion of genomic markers, we expect that a limited number of PCs explains most of the genetic variation, even in a polygenic trait. In fact, experience from animal breeding indicates that very good genomic prediction accuracy can be obtained by using a subset of PCs, usually a number corresponding to about 50-70% of individuals in a core sample that is representative for the genetic diversity in the population (Ødegård et al., 2018). However, it is not clear to what degree this applies to wild systems, with typically higher effective populations sizes than the heavily inbred domestic populations, which implies a larger number of independently segregating chromosome segments (*M_e_*) and thus lower expected LD between the SNP markers and quantitative trait loci (QTLs) than in animal breeding. The actual choice for the number of PCs *k* is therefore not trivial and can vary greatly from case to case (Dadousis et al., 2014). Simply speaking, we need to find a balance between over- and under-fitting to accommodate for the bias-variance trade-off (James et al., 2013). One way to avoid over-fitting is by combining PC regression with ridge-based shrinkage, irrespective of *k*, as described in the following.

#### 2.2.2 Bayesian Principal Component Ridge Regression

Simplified versions of model (3) have previously been tackled via frequentist methods, either by using PC regression based on a standard or a partial least squares approach (Solberg et al., 2009; Dadousis et al., 2014), or later via PC ridge regression (PCRR, Ødegård et al., 2018). However, those applications were concerned with data from animal breeding and did not need to consider random (environmental) effects to account for the complexity of wild systems. More specifically, while we usually want to fit the full model (3), previous modeling attempts ignored the **Wd** component, and sometimes even the fixed effects **Xb**. In order to reach full flexibility in our model formulation, we here combine the idea of PC ridge regression (Ødegård et al., 2018) with a Bayesian approach based on INLA (Rue et al., 2009).

A previous application from plant breeding indicates that INLA is a suitable and promising candidate for our purpose (Selle et al., 2019). Here we expand the considerations from Selle et al. (2019) by discussing how to find suitable priors that impose an appropriate level of shrinkage. To this end, we employ an approach where the eigenvalues are used as scaling factors for the variances of PC-effects (Macciotta et al., 2010), which is equivalent to letting the columns of **Z★** have a variance proportional to the corresponding eigenvalue of the SNP covariance matrix. To ensure numerical stability (and without changing the generality of the results), we can then scale (*i.e.*, divide) the variances of all PCs by the standard deviation of the first PC, so that var(PC*_i_*) *≤* 1 for all PCs. We then set a 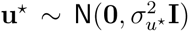 prior on the PC-effects, such that the PCs are identically distributed, with a suitable prior for 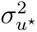 (which we will discuss below). This, together with the scaled PCs, implicitly assumes that the variance explained by a PC is monotonously related to the variance it explains in the breeding value of the trait of interest, which seems plausible for a highly polygenic trait under the infinitesimal model assumption (see Figure S3). In contrast to the marker-based Bayesian model, where the choice of the prior on **u** is critical due to *m ≫ N*, we have the advantage of fewer variables than data points (*k < N*) and priors thus become less influential (Gianola, 2013).

The Bayesian formulation of model (3) combined with the 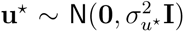 prior results in a model that we denote as Bayesian PC ridge regression (BPCRR). In fact, the normal prior on **u★** chosen here corresponds to a (likelihood-based) ridge shrinkage factor 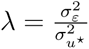 (*e.g.*, Fahrmeir et al., 2004), which also appears in the SNP-based GBLUP estimator (Gianola, 2013; Ødegård et al., 2018). The model, including any additional fixed and random effects, can be fitted in `R-INLA` in one step. The ridge-regression type of shrinkage is thereby only explicitly imposed on the genomic components, while the priors for the remaining parameters can be suitably chosen according to convenience and/or prior knowledge (*e.g.*, Wang et al., 2018, Chapter 5.4.1).

#### 2.2.3 Priors for the PC-effects

What remains is to assign a suitable prior to the variance 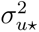 of the PC effects. To this end, let us first look at the corresponding ridge (GBLUP) version of model (2). In this case, an appropriate level of shrinkage is imposed via the shrinkage factor 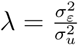 with SNP-effect variance 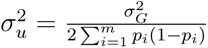, where *p_i_* is the allele frequency at locus *i*, meaning that the denominator is the SNP variance summed over all *m* loci (Gianola, 2013; Ødegård et al., 2018). However, this choice of the denominator for *σ*^2^ is assuming independence between the SNPs, which is problematic due to both LD and the family structure within populations (*e.g.*, Charmantier et al., 2014; Uffelmann et al., 2021). The independence assumption is, in contrast, automatically fulfilled when we go from the marker-based model (2) to the PC-based regression model (3). In order to ensure a corresponding level of ridge-type shrinkage, we therefore adopt the idea to find the corresponding value for the PC effects variance by scaling the additive genetic variance by the sum of the variances over the *k* independent PCs that are included in the model

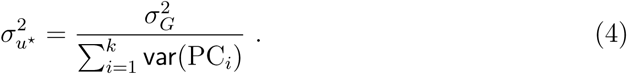

Thanks to the above mentioned scaling of the PC variances to var(PC_1_) = 1 and var(PC*_j_*) *<* 1 for all *j ≥* 2, the denominator in (4) is always smaller than *k* and thus remains numerically stable in the implementation. Note that the idea to leave PC-variances proportional to the variance they explain in the data implies that PCs with larger variance will be subject to less shrinkage of the corresponding PC-effect, which is an often overlooked feature of ridge regression (Gianola, 2013), and essentially (again) reflects that PCs that explain more variance in the data should also have the opportunity to explain more variance in the response. Note that this scaling of the PCs is in contrast to the common ridge regression scaling, where all variables (*i.e.*, here all PCs) would be standardized to have a variance of 1, which would correspond to 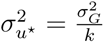

An obvious implication of setting 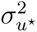 to the value given in (4) is that we need to know *σ_G_*^2^, ideally without uncertainty. Even though popular existing methods for genomic prediction like BayesR rely on similar prior knowledge (Gianola, 2013), the requirement is somewhat circular if the aim is to actually *estimate σ_G_*^2^. Even though it is typically possible to obtain reasonable prior guesses for *σ_G_*^2^, for example from analyzing a smaller subset or from a previous study, we actually do not need to impose such a “point prior” on 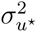, but can instead afford to give it a relatively uninformative hyper-prior thanks to the fact that we are in the *k < N* regime. To underline this point, we will do a sensitivity analysis by carrying out all our analyses for both fixed 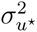 priors suggested in (4), using a “good prior guess” from earlier analyses, as well as using the Gamma prior 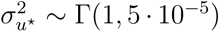, parametrised with shape and rate, which corresponds to the very naive default prior in the `R-INLA` framework. Even though the respective Gamma prior has been criticized in other contexts (Gelman, 2006; Hodges, 2013), its use led to quite accurate results in a related study (Selle et al., 2019). Users may of course choose other priors, for example a distribution around the value in (4) with varying degree of uncertainty.

#### 2.2.4 Optimal number of PCs for genomic prediction

Recall that the quantitative genetic statistical models discussed here have two main purposes: estimation of *σ_G_*^2^ and the prediction of breeding values. In the former case, the estimated *σ_G_*^2^ is expected to monotonically increase for an increasing number of PCs (*k*), where the respective curve asymptotically flattens out once a substantial amount of variance is explained. On the other hand, if we want to predict breeding values in samples that have not been used to fit the model, careful evaluation of the trade-off between bias and precision is required, and only an intermediate number of PCs should be used in the modeling procedure (Solberg et al., 2009; Dadousis et al., 2014). In principle, the optimal number for *k* can be found by fitting many models for a dense grid of different numbers of PCs and then choosing the one with highest accuracy. However, this approach renders the overall procedure inefficient due to the need to repeatedly fit the models for various numbers of *k* (Solberg et al., 2009). Here, we therefore employ theory from animal breeding and human genomics, where a heuristic formula for expected prediction accuracy has been derived as a function of sample size (*N*), the number of independent components with estimated effects (*M_e_*) – usually the number of independent SNP effects – as well as the proportion of variance explained by those components (*h_M_*^2^), the SNP-based heritability for the *M* SNPs that are included in a particular model

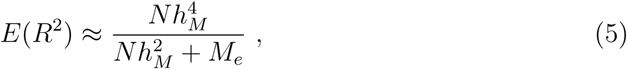

(Daetwyler et al., 2008; Wray et al., 2013, 2019). *R*^2^ stands for the proportion of phenotypic variance explained in the out-of-sample prediction, which is directly related to the expected prediction accuracy (*i.e.*, the expected correlation between the predicted breeding value and the phenotype, see Section 2.3.2). However, since we operate with *k* PCs instead of *m* SNPs, we modify equation (5) such that *M_e_* (which typically is *< m*) is replaced by *k*, where *h_k_*^2^ denotes the proportion of variance explained in the phenotype by the respective number of PCs

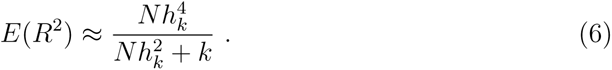

The rationale is that the *M_e_* SNP effects are assumed independent, which the *k* PCs fulfill by construction. We thus need to find an approximation for *h_k_*^2^, but since we in general (again) want to circumvent the estimation of this quantity for a grid of *k* due to efficiency reasons, we further assume that *h_k_*^2^ is proportional to the variance that the respective PCs explain in the SNPs. This proportionality assumption is plausible for the infinitesimal model, and Figure S3 in the Appendix illustrates its approximate validity for the cases studied here. For a given *k*, we therefore approximate

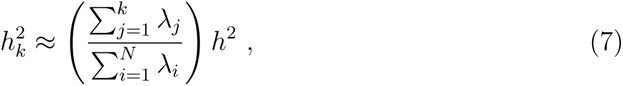

where *h*^2^ is the heritability of the trait of interest, which we either assume to be approximately known (or otherwise close to 0.3, see Hansen and Pélabon, 2021). Each eigenvalue *λ_j_* of the SVD corresponds to the SNP-variance explained by the respective PC, thus *h_k_*^2^ corresponds to *h*^2^ multiplied by the proportion of variance explained by the first *k* PCs. For a given set of SNPs (and their corresponding SVD), the optimal *k* thus only depends on the sample size and the heritability of the trait, given that the imposed assumptions are approximately correct.

A final important consideration is that the BPCRR approach imposes shrinkage on the PC effects, with stronger shrinkage on PCs that explain less variance (Macciotta et al., 2010; Gianola, 2013). For the scaling of PC variances used here, we therefore do not expect a significant decrease in prediction accuracy when adding a larger number of PCs than the estimated “best” *k* from equation (6). As a consequence, *k* that maximizes prediction accuracy in (6) should be seen as a lower limit, and not necessarily as the final choice. We will come back to this point later.

### 2.3 The house sparrow application

#### 2.3.1 The data set

All methods were tested and validated with empirical data from an individual-based long-term study on house sparrows in an archipelago at the coast of northern Norway (Baalsrud et al., 2014). We used data on adult house sparrows present on eight islands in the study system during the breeding seasons (May-August) 1998-2012 (Niskanen et al., 2020). Birds were marked with a numbered metal ring and a unique combination of plastic color rings for later individual identification. Some individuals were ringed in the nest and only observed as adults, whereas others were captured with mist nets as adults and then measured for morphological traits such as tarsus length (to nearest 0.1 mm), wing length (to nearest mm) and body mass (to nearest 0.1 g). Because house sparrows are sedentary and only ca. 22% perform short-distance natal dispersal, some adult individuals were captured and measured multiple times during their lives, either on the island they were born or on a neighboring island (Saatoglu et al., 2021). From all ringed individuals a small blood sample (ca. 25 *µL*) was taken to obtain DNA (Lundregan et al., 2018; Niskanen et al., 2020), which was the basis for high-throughput individual genotyping of SNP markers distributed across most chromosomes in the house sparrow genome by using a custom Axiom 200K SNP array (Lundregan et al., 2018). After quality control, genotype data on 182 848 polymorphic high-quality SNPs were available for 3032 adult individuals (Lundregan et al., 2018; Niskanen et al., 2020). Less than 0.6% of all SNP-genotypes were missing, and those were mode-imputed.

#### 2.3.2 Statistical modeling

We estimated *σ_G_*^2^ and assessed the accuracy of genomic prediction in the house sparrows based on the three continuous traits body mass, tarsus length and wing length.

Among the genotyped individuals, we had phenotypic measurements for 1918, 1915 and 1912 individuals, on these traits, respectively. About half the individuals had only one measurement for each trait, roughly 25% had two, and the remaining individuals had three or more (maximum 13) measurements. The total number of measurements were 4249 for body mass, 4368 for wing length and 4373 for tarsus length. Our model included as fixed effects sex (1 for male, 0 for female), the genomic inbreeding coefficient *F_GRM_* (Niskanen et al., 2020), age at the time of measurement (in years), and the month when the measurement was performed (May to August, denoted 5, 6, 7 and 8), as well as variables reflecting proportional genetic origin from one of three genetic groups (inner, outer and other islands), mirroring the habitat structure of the study meta-population (see Muff et al., 2019, for details on how the genetic group variable was derived). The permanent individual effect, the island where the individual was measured, and the year of the measurement were modeled as additional Gaussian independent random effects.

##### Estimation of additive genetic variance

To investigate the estimation of *σ_G_*^2^ using the BPCRR method based on model (3), we used both a fixed prior for the PC effects variance 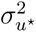 as given in (4) with a trait-specific prior estimate of *σ_G_*^2^ (denoted from now on as *BPCRR fixed*), as well as for the vague default 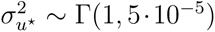 hyper-prior in INLA (denoted as *BPCRR default*), as described in Section 2.2.3. Models were fitted with different numbers of PCs starting with *k* = 100 and increasing in steps of 100 to obtain a grid of *k*-values. The expectation is that the estimated *σ_G_*^2^ steadily increases with *k*, but that the the curve flattens out when most of the genetic variance in a given trait is explained by the PCs. For comparison, we also derived posterior distributions of *σ_G_*^2^ using the genomic animal model and BayesR. In all cases, except BayesR, we used `R-INLA` to derive the marginal posterior distributions, where we assigned independent N(0, 10^4^) priors to all fixed effects and and penalized complexity priors (Simpson et al., 2017) PC(1, 0.05) to the variances of the remaining random effects (*i.e.*, for all except 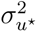). For BayesR, we used the R package hibayes (Yin et al., 2022) with data-inferred hyper-parameters for the inverse Chi-square distribution that is assumed for the marker effects, as described in Yin et al. (2022), although the results are expected to be insensitive towards the actual choice of the prior (Ashraf et al., 2022). Further computational details are given in Section 2.5. All methods were compared with respect to computation time.

##### Genomic prediction

We evaluated the prediction accuracy of BPCRR for the number of PCs found by optimizing formula (6), as well as for *k* = 50, 100, 200, 500, 1000, 1500 and 1900, both for fixed 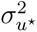 and `R-INLA` default priors, via a 10-fold cross-validation (CV). Note that folds were defined by selecting individuals and not single measurements, to avoid having individuals appear in several folds. Here, prediction accuracy was reported as the correlation between the predicted breeding values *ĝ* and the mean phenotype *y* over all the repeated measurements per individual cor(*ĝ, ȳ*). Note that this is related to the predicted accuracy *R* from formula (5), which corresponds to cor(*ȳ, y*) = cor(*ĝ, y*) *· h*, but the correspondence is not exact in our case because we replace *y* by *ȳ*.

The results were benchmarked against the GBLUPs from the genomic animal model and predictions from BayesR, both in terms of prediction accuracy and computation time. Finally, we assessed the bias of the genomic prediction results by regressing the mean observed phenotype per individual against the predicted breeding values and reported the actual slopes *β*_1_. If the estimates are unbiased, the expected value for *β*_1_ corresponds to 1.0 (Meuwissen et al., 2001).

### 2.4 Simulation

Based on the set of existing SNPs from the house sparrow meta-population, we carried out a simulation study where we generated phenotypes according to model (2), but without any fixed or random effects (**Xb** = **Wd** = 0). Assuming a genetic architecture that resembles the one found for body mass for the house sparrow data, we sampled the SNP effects (*u_j_*) for a hypothetical polygenic trait from a mixture of four zeromean normal distributions as

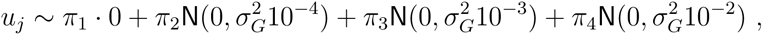

with ***π****^⊤^* = (0.95, 0.04, 0.008, 0.002), indicating that each effect *u_j_* stems from either of the four distributions with the probabilities given in the ***π*** vector. Using the respective set of markers *u*_1_*, …, u_m_*, we then generated a set of breeding values as a linear combination *g_i_* = Σ*_j_ x_ij_ · u_j_*, where *x_ij_* denotes SNP *j* for individual *i*. The set of *g_i_* were then scaled such that *σ_G_*^2^ = 0.33, and phenotypes were generated as *y_i_* = *g_i_* + *ε_i_* with *ε_i_ ∼* N(0*, σ_ε_*^2^) using *σ_ε_*^2^ = 0.67, in order to attain a heritability of *h*^2^ = 0.33. For both the estimation of *σ_G_*^2^ and genomic prediction, we carried out the same comparisons and benchmarking as for the analysis of the empirical house sparrow example. Note that, since the simulated data actually are generated according to the underlying modeling assumptions of BayesR (Erbe et al., 2012; Gianola, 2013), that approach has an advantage compared to the other models.

### 2.5 Computational details and software

All analyses for the genomic animal model and BPCRR were performed with version 20.3.17 of the the `R-INLA` R package (Martins et al., 2013; Rue et al., 2017) in the statistical software R (R Core Team, 2021) version 4.0.3. The GRM for the genomic animal model was derived using the van Raden-method (VanRaden et al., 2009) implemented in the AGHmatrix package version 2.0.4 (Rampazo Amadeu et al., 2016). The SVDs needed in BPCRR were done in PLINK (Chang et al., 2015), although (less efficient) ways to do the SVD are possible directly in R, for example using the package RSpectra (Qiu and Mei, 2019). In applications where not all individuals had measured phenotypes, the SVD was done for the respective sub-matrix including SNP data only from the phenotyped individuals that were included in the model, since the inclusion of additional individuals in the SVD calculation would otherwise introduce unwanted and unnecessary noise into the PCs. For the BayesR analyses we used the package hibayes (Yin et al., 2022) version 3.0.0 in the R-version 4.2.1. The length of the burn-in and total number of iterations for the MCMC chains in hibayes were selected for each case according to visual inspection of the convergence plots, in order to ensure a good trade-off between accuracy and computational time. The thinning interval was set to 10. Unless stated otherwise, all analyses were performed on a local high-performance computing cluster (Själander et al., 2019).

## 3 Results

### 3.1 Estimation of additive genetic variance

As expected, the estimated *σ_G_*^2^ for the three traits body mass, tarsus length and wing length from the house sparrow example, as well as for the simulated case, increased with an increasing number of PCs included in BPCRR (Figure 1). In all cases, the increase in estimated *σ_G_*^2^ asymptotically flattened out and converged towards the final value, but this happened in different ranges for the different traits and depended on the actual prior used in BPCRR (Figure 1). As expected, when using `R-INLA`’s default prior for BPCRR, more PCs were needed for convergence and the 95% credible intervals (CIs) were a bit wider than for the informative (fixed) prior derived in (4). For fixed prior variance on 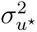, 1000 PCs typically resulted in good approximations for *σ_G_*^2^, whereas more PCs were needed when uninformative priors were used.

**Figure 1:**
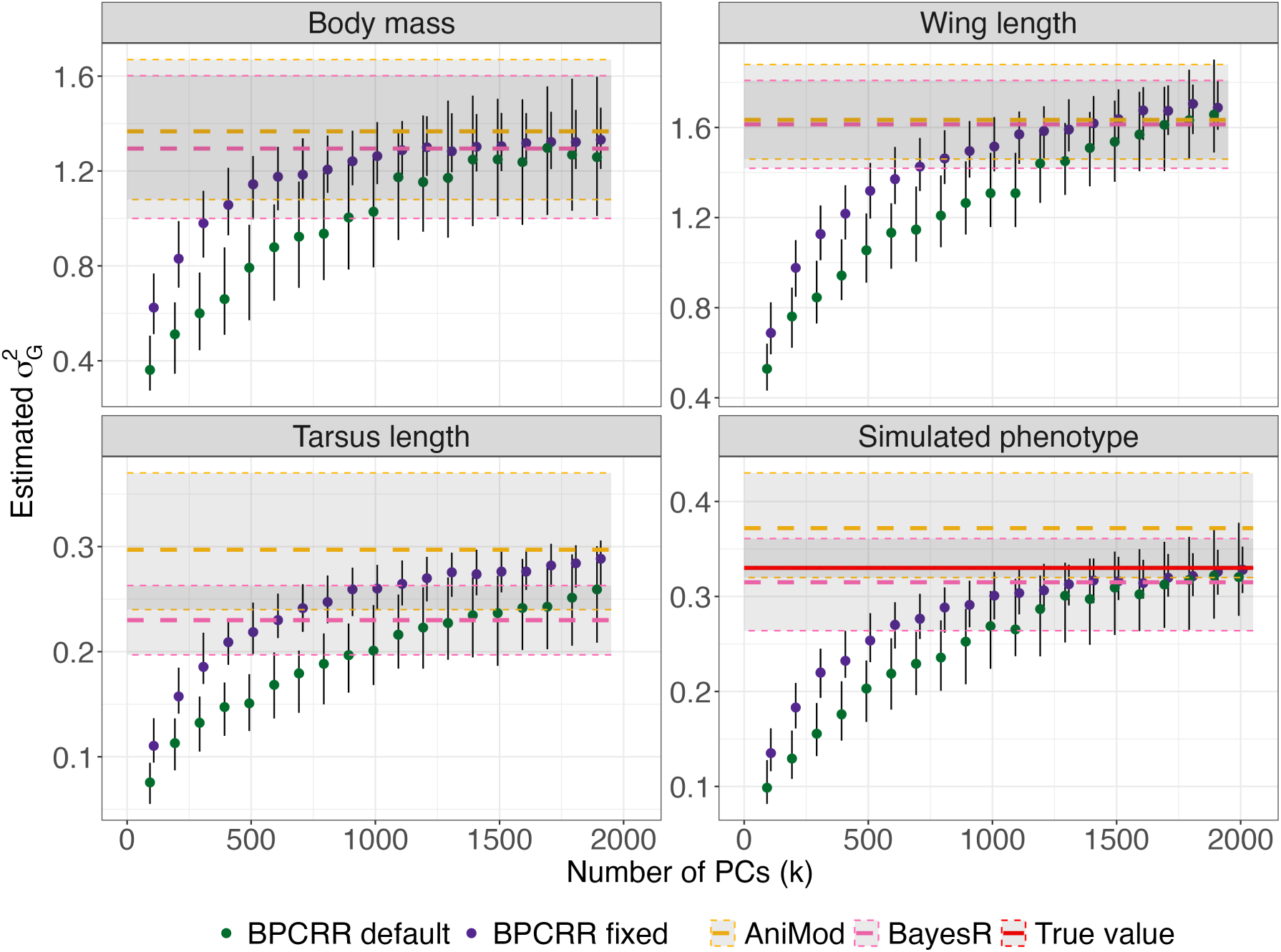
Estimates of *σ_G_*^2^ using BPCRR dependent on the numbers of PCs (with fixed and INLA default priors, in green and blue, respectively), BayesR (in pink) and the genomic animal model (AniMod, in orange) for the three traits from the house sparrow example, as well as for the simulated example. Error bars and shaded areas correspond to 95% credible intervals. The solid red line corresponds to the true value for the simulated phenotype.

The results of BPCRR, BayesR and the genomic animal model are generally in good agreement. For tarsus length, however, BayesR predicted a somewhat smaller and the animal model a larger *σ_G_*^2^ compared to the (converged) BPCRR results. In addition, the genomic animal model seemed to overestimate *σ_G_*^2^ in the simulation study, suggesting that the method is sensitive to the violation of the assumption that the marker effects stem from one normal distribution, instead of the mixture that was used to generate the data.

### 3.2 Genomic prediction

#### 3.2.1 Selecting the number of PCs in BPCRR

By using equation (6) and the approximation for *h_k_*^2^ from (7), we generated a curve of the expected prediction accuracy *E*(*R*^2^) against the number of PCs *k* (Figure 2). For approximation (7) we used the *h_k_*^2^-estimates from the genomic animal models fitted to the sparrow example, namely *h*^2^ = 0.28 for body mass, 0.29 for wing length and 0.47 for tarsus length, while the true value *h_k_*^2^ = 0.33 could be used for the simulated phenotype. Sample sizes were 1918, 1915 and 1912 for body mass, wing length and tarsus length, and *N* = 3032 for the simulation, respectively. Using those numbers, the *k* that optimized the theoretical prediction accuracy for the different cases was 527, 541, and 688 for body mass, wing length and tarsus length, respectively, and 728 for the simulated phenotype. By adding boxplots reflecting the observed prediction accuracy from a 10-fold CV using the BPCRR procedure with the respective number of PCs and fixed prior variance for 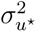, we found that there typically is a relatively broad range of values that leads to similarly good results. As mentioned in Section 2.2.4, the ridge shrinkage induced on the PC effects ensures that prediction accuracy remains high even when *k* larger than the “best” estimate from equation (6) are used. This finding is reassuring, since we might often not know the true *h*^2^ exactly, and the quality of approximations taken in formulas (6) and (7) may vary from case to case. Importantly, this result still holds when the fixed variance 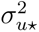 was replaced by INLA’s default prior (Figure S4), which is relevant for cases where we do not have much preliminary knowledge about *σ_G_*^2^ and *h_k_*^2^.

**Figure 2:**
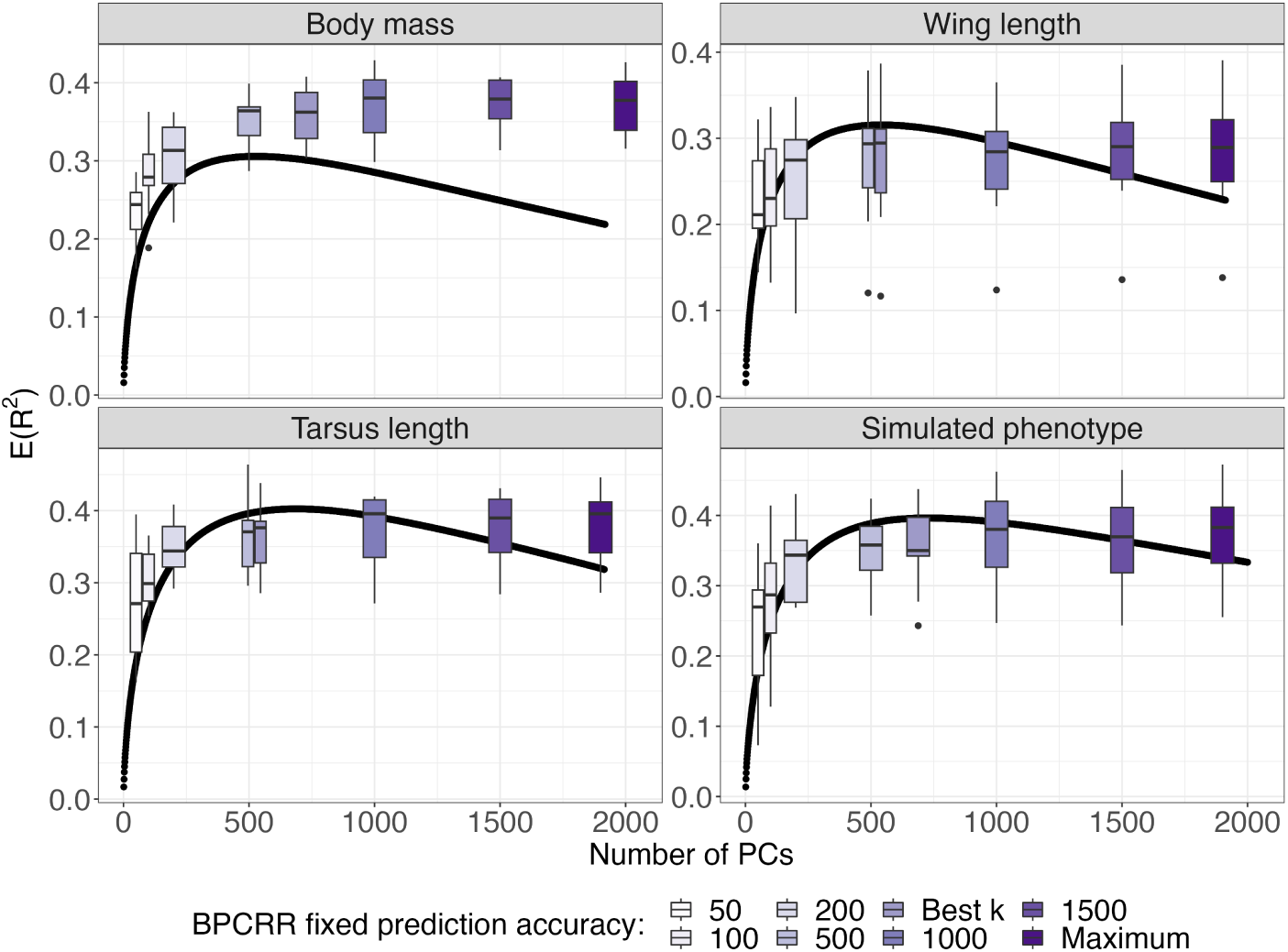
Expected prediction accuracy *E*(*R*^2^) using formulas (6) and (7), in dependence of the number of PCs (*k*) and for *h*^2^ and *N* values that correspond to the respective cases (black line). The actually observed prediction accuracies from 10 cross-validation runs fitted with fixed ridge priors on the PC effects with 8 different *k*s ranging from 50 to 1900 (including the “best *k*”) are added as boxplots for comparison.

#### 3.2.2 Assessing the quality of genomic prediction

The prediction accuracy for BPCRR was consistently high and typically equally good or slightly better than the GBLUPs from the genomic animal model and for BayesR, irrespective of whether default or fixed priors were used (Figure 3, top). In line with the results found in Section 3.2.1, the results for BPCRR suggest that it typically is safe to pick a slightly larger number of PCs than the value that optimizes equation (6), since all values between the optimal number and maximum number of PCs gave satisfying prediction accuracy. In our analysis, values for *k* between 500 and 1000 seemed to be a robust choice that gave a stable trade-off between computational efficiency and accuracy (Figures 3 and 4). Interestingly, the prediction accuracy for tarsus length was considerable lower for the GBLUPs compared to both BPCRR and BayesR, indicating that the assumptions for the genomic animal model that generated the GBLUPs might have been violated, potentially due to a more oligogenic architecture of the trait, as found in leg bone traits in other species (*e.g.*, Silva et al., 2017; Ashraf et al., 2022).

**Figure 3:**
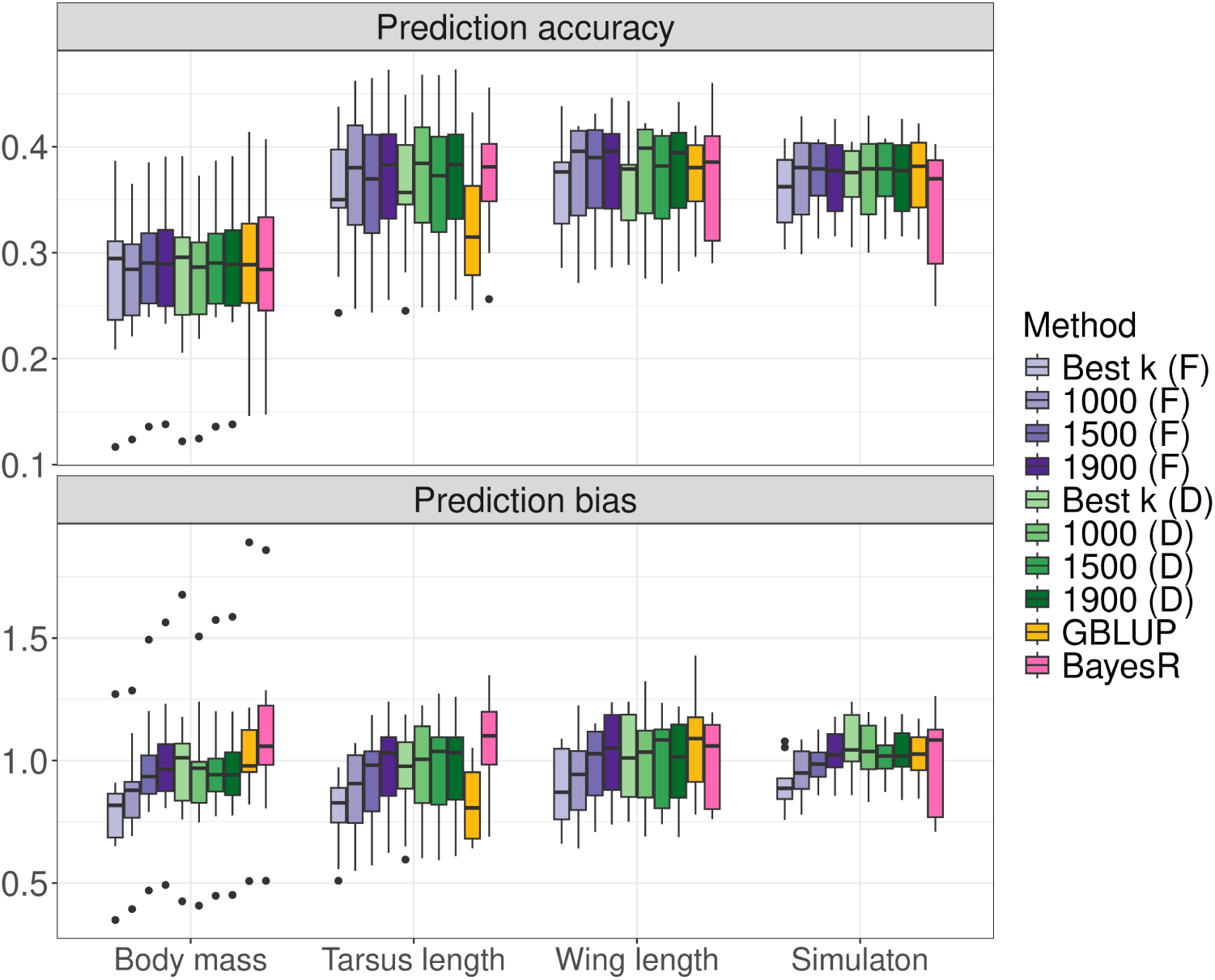
Assessment of prediction accuracy (top) and prediction bias (bottom) for BPCRR for a selection of the numbers of PCs shown in Figure 2, with *fixed* (F, in blue shades) and *default* priors (D, in green shades), as well as comparisons to the GBLUPs (orange) from the genomic animal model and for BayesR (pink). The prediction accuracy is defined as the Pearson correlation between predicted breeding values *ĝ_i_* and mean phenotypes 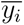 per individual. Prediction bias is assessed via the distribution of regression coefficients when regressing all 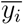 on *ĝ_i_*. The *y*-axes were cut at 0.5 and 2.0, respectively, where one case with high prediction accuracy for the GBLUPs for wing length, and two cases with high prediction bias for the simulation case for BayesR were cut for better visibility.

**Figure 4:**
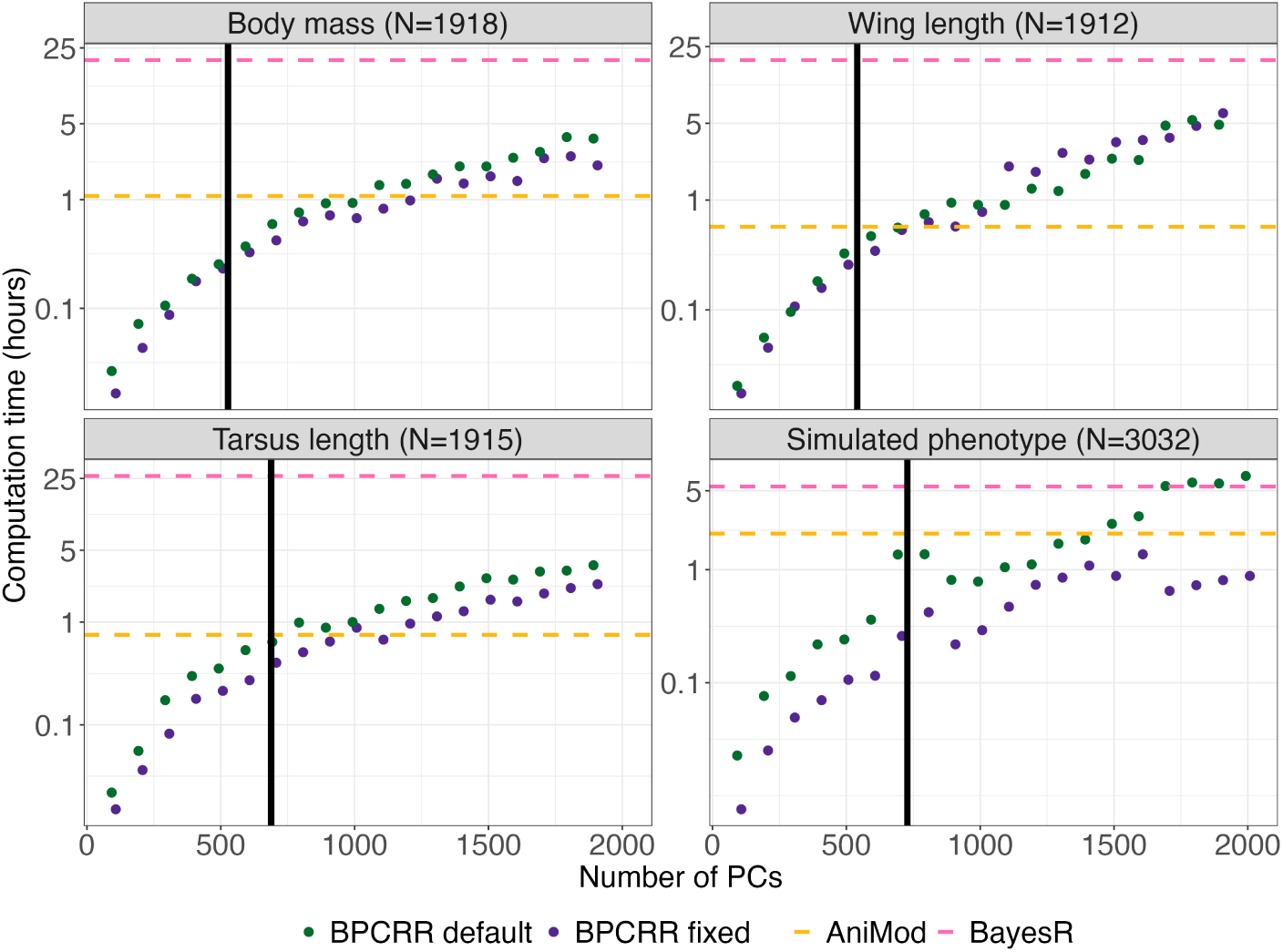
Computation times for BPCRR (fixed and INLA default priors, in green and blue, respectively), BayesR (in pink) and the genomic animal model (AniMod, in orange). The black vertical lines indicate the number of PCs obtained from maximizing equation (6). Note that the *y*-axis is on log-scale, thus a linear increase in the graph corresponds to an exponential increase in absolute values.

In terms of bias, the slope coefficients *β*_1_ for the regression of mean observed phenotypes *y_i_* against *ĝ_i_* did not reveal any apparent deviations from 1.0 when default priors were used in BPCRR (Figure 3, bottom). When using the shrinkage prior from equation (4), on the other hand, the actual values of the breeding values were somewhat underestimated unless almost all PCs were included, most likely due to the shrinkage induced by the prior. Finally, both the GBLUPs and BayesR were unbiased for body mass and wing length, as well as for the simulation, but showed a downward and upward bias, respectively, in the case of tarsus length. An interesting observation is that using the default priors in INLA led to both high accuracy and low bias even for relatively small number of PCs. The major benefit of using the fixed priors thus lays in the gain of computational efficiency, as we will see in the next section.

### 3.3 Efficiency of computations

Computation times were compared for procedures ran on the same high-performance cluster as all other analyses (Själander et al., 2019). Even though, in contrast to hibayes, `R-INLA` is running in parallel by default, we always assigned exactly one core to fit each individual model used in this comparison for conservative benchmarking. Note that the cluster’s queue system automatically assigns CPUs to the jobs, whereas not all the CPUs have the exact same hardware. Despite this, we see a clear pattern of increasing computation times with a growing number of PCs (*k*) for the BPCRR method (Figure 4), and that the method is faster than fitting the genomic animal model for values of *k* that are obtained from maximizing equation (6) for the three traits and the simulation. For *k* = 1000, computation times for BPCRR and the genomic animal model are comparable for the three traits, and considerably lower for BPCRR for the simulation. The last observation is in line with the fact that the computational benefits are most pronounced for the simulation, namely because the latter has larger *N* than the three traits form the house sparrow example, and thus the computational benefit of the BPCRR becomes more apparent. The BPCRR method was most efficient when the shrinkage prior (4) was used, and slower for the INLA default.

The BayesR implementation via the hibayes R package used here, which was chosen in order to handle the full model including all the fixed and random effects like the other two methods, was clearly the most expensive in terms of computation time. Model complexity and the choice of the trait strongly affected the actual number of MCMC iterations needed until convergence (Figure S5). While convergence observed from the MCMC samples occurred relatively quickly for the simulated phenotype, wing length and body mass, more samples were needed in the case of tarsus length, as indicated by the convergence plots observed from MCMC chains that were ran on the full datasets for long enough until we observed convergence (Figure S5). Based on those long MCMC chains, we then selected a fair trade-off between accuracy and computation time for all the following analyses done here. For body mass and wing length, we then chose a burn-in of 25 000 and a sample size of 50 000, for tarsus length a burn-in of 100 000 and 50 000 samples, and finally a burn-in of 10 000 and a sample size of 20 000 for the simulation study, which, unsurprisingly, was the least problematic in terms of convergence. Importantly, we would like to stress that the computational burden is similar for the commonly used implementation of the BayesR procedure (Moser et al., 2015; Ashraf et al., 2022; Hunter et al., 2022), which is based on the two-step approach mentioned in Section 2.1.3 (see Appendix A).

## 4 Discussion

We have introduced an approach for approximate estimation and prediction of key evolutionary parameters in wild systems using a Bayesian principle-component regression framework that we combined with the idea of shrinkage priors. The proposed method is suitable to efficiently fit Bayesian mixed effects models with an arbitrary number of fixed and random effects, in addition to an additive genetic component derived from genomic data. The ability to handle such models is crucial to account for the typically heterogeneous environmental conditions, population structure and effects like inbreeding, sex or age that are relevant to suitably model phenotypic data of wild study systems. The results indicate that the BPCRR method is useful for both the estimation of additive genetic variance and the prediction of genomic values, although its major strength in terms of computational efficiency gains clearly lays in its application to genomic prediction. The method was tested on a real application with empirical data from a system of house sparrows, as well as with a simulation study where we used the house sparrow SNPs as a basis to generate breeding values for a hypothetical trait.

As a main result, we found that BPCRR gives high prediction accuracy even when only 25-50% of all PCs are included. This result is independent of whether shrinkage priors (4) or naive Gamma priors 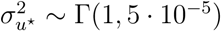 are used for the PC-effect variance. In the latter case, however, computation in `R-INLA` is slower. Since the choice of sensible priors often considerably impacts the computational efficiency of INLA, we expect to see an even larger difference between computation times in these two cases when the sample size *N* increases. On the other hand, the absence of shrinkage in the Γ(1, 5 *·* 10*^−^*^5^) prior ensures unbiased prediction already for small number of components (Figure 3). In any case, we expect the efficiency benefit for BPCRR compared to the genomic animal model/GBLUPs or BayesR to increase with increasing number of individuals *N* and/or markers *m*. Importantly, when *N* within a homogeneous population increases, we do not expect that considerable more PCs are needed to obtain good predictions (see formula (6) and how *N* affects *E*(*R*^2^)). A trend for this effect is reflected in Figure 4: The simulation has largest *N* among the four investigated data sets, and it is the case where differences in computational efficiency between BPCRR and the other methods become somewhat more pronounced, especially in comparison to the genomic animal model.

Here we have chosen an implementation of BayesR based on the hibayes R package (Yin et al., 2022), which is able to handle the full model (2), in order to have a one-to-one comparison to the BPCRR and genomic animal model methods. On the other hand, genomic prediction based on BayesR is usually done in a two-step manner, where the residuals or the random individual-specific (ID) effects from a pre-fitted linear mixed model (LMM) that includes all fixed and random variables except the genetic effect, are used as the target response in BayesR. However, a potentially under-recognized problem is that the *V_A_* may be heavily underestimated in this case, in particular for traits that have large residual variation (Figure S1). Own preliminary results indicate that a remedy can be to instead use the sum of the ID effect and the mean of the residuals over the repeats of an individual as the new response in the two-step procedure leads to correct *V_A_* (Figure S1). In our understanding, the reason why we see the described effect is that, due to the correlation structure defined by the ID effect in an LMM, the respective ID estimate is partially confounded with the residual, thus the latter absorbs part of the breeding value. The missing signal can only be extracted when adding the (mean) residual to the ID effect in the new response. However, further investigations are needed to obtain a better understanding of this aspect, since it is not the scope of this work.

In any case, there is no doubt that environmental factors and the relevant fixed effects need to be accounted for in the statistical modeling procedure in one way or another. A crucial advantage of integrated procedures that fit the full model in one step is that uncertainties in parameter estimates can be properly accounted for at all levels of the analysis. In addition – and maybe even more importantly – a full model can account for potential interactions between effects that otherwise are included in two distinct steps. This is particularly relevant if we wish to study *G × E* interactions, which have been shown to be relevant in essentially all domains, such as for humans, wild and bred animals and plants (Ackermann et al., 2001; Jarquín et al., 2017; Nguyen et al., 2017; Wang et al., 2019; Christensen et al., 2021; Jarquín et al., 2021; Di Leo et al., 2022). However, modeling such interactions is challenging, and it is especially not straightforward with a two-step approach.

Sample sizes of individuals with high-density genomic data for wild animal populations are expected to increase in the coming years. Even though one of the main strengths of BPCRR is its scalability, performing the SVD for very large *N* and/or high *m* can become computationally challenging. Animal breeders, who often work with hundred thousands of individuals, have proposed two complementary strategies that may be adopted in this case: performing chromosome-wise SVDs that yield chromosome-wise PCs, and deriving the SVD for a limited “core sample” of individuals that reflects the major part of the genetic variation in a population of interest (Ødegård et al., 2018). The first idea can easily be adapted to wild systems if chromosome-level genome assemblies are available, as it would then be relatively straightforward to perform the SVD chromosome-wise and select the set of PCs that explains a pre-defined minimum amount of chromosome-specific variance (typically 99%). This may lead to an interesting partition of additive genetic variance into shares explained by each chromosome. It is, however, less clear whether the selection of a representative core sample of individuals is possible and suitable, since we are not in the same regime as the typically relatively homogeneous breeding systems. For now, we did not consider any of those strategies, but keep them in mind as options in future applications.

Previous studies indicate that the performance of genomic prediction methods depends on the genetic architecture of the analyzed trait (*e.g.*, Moser et al., 2015; Meher et al., 2022), and it remains to be investigated under which circumstances BPCRR gives the largest benefit. By construction, we expect BPCRR to be particularly suitable for complex traits with many small-effect loci, where the amount of variance explained in the phenotype approximately scales linearly with the proportion of variance explained by the PCs that are included in the model (see again Figure S3). Interestingly, while the genomic animal model works best in the regime of the infinitesimal modeling assumption where we assume that the original marker effect sizes **u** all stem from the same distribution (Meuwissen et al., 2016), BPCRR does not impose such a restriction. In fact, the distribution of the marker-specific effect sizes, which may be possible to obtain by back-transforming the PC-effects (Ødegård et al., 2018), heavily depends on the loadings in the individual PCs. It is thus not surprising that BayesR and BPCRR show similar prediction accuracy for tarsus length (Figure 3), which is a skeletal trait that potentially is less polygenic than body mass and wing length (Silva et al., 2017; Hansson et al., 2018; Yengo et al., 2022), while the genomic animal model generates GBLUPs that are somewhat less precise in this case. A similar observation was made for the two skeletal traits “foreleg” and “horn length” in a study of wild Soay sheep (Ashraf et al., 2022). However, BPCRR still relies on the assumption that the variance explained in the SNPs is approximately proportional to the variance in the response, thus the method is not constructed to analyse Mendelian traits, for example.

In our comparison we chose BayesR as a reference due to its many appealing features, high efficiency and accuracy. A very useful feature of BayesR is that it allows to investigate the trait architecture, and in particular the genetic variance explained by each marker (Erbe et al., 2012; Moser et al., 2015). As mentioned above, if interest centers around finding marker-specific effects, deriving them post-hoc from the BPCRR results may be possible. Here, however, we were interested in the suitability of the method with respect to *V_A_* estimation and genomic prediction, thus we limited our investigations to those aspects. We conclude that BayesR and BPCRR are comparable in terms of prediction accuracy, with the latter having the major advantage that the full model can be fitted in one step and without the need for time-consuming MCMC sampling. Moreover, BPCRR is flexible, and almost arbitrary modeling extensions are possible, such as adding further partitions of the total genetic variance into additive and dominance components, or accounting for *G × E* interactions. On the other hand, neither BayesR nor BPCRR are likely to be the first choice when the main goal is to estimate *V_A_*. In fact, both methods partially depend on prior empirical knowledge about *V_A_* (Gianola, 2013), even though the uninformative Gamma default priors provided by `R-INLA` that do not rely on such knowledge converged towards correct *V_A_* estimates in BPCRR. The low efficiency in this latter case may however render default priors unfeasible for larger data sets, and informative shrinkage priors that rely on preliminary knowledge on *V_A_* are then anyway required.

One crucial aspect that renders BPCRR competitive for genomic prediction is that we can use formula (6) to find an appropriate number of PCs *k*. Due to the shrinkage imposed on the PC effects, formula finds a lower bound for *k*, and prediction accuracy remains robustly large for values *≥ k*. Therefore, choosing a slightly larger number than the optimum from (6) seems to be a safe choice, while too large numbers would unnecessarily increase the computational burden. It is also important to keep in mind that formula (5), which forms the basis for our arguments, relies on many underlying assumptions that might be violated. The formula does, for example, not consider that our models contain fixed and random effects apart from the genomic value, and it assumes that there is no structure in the population (Wray et al., 2013, 2019). We thus expect to find better approximations when the formula is properly adjusted, but finding such improvements is an active area of research (Dekkers et al., 2021).

The rapidly increasing volumes of genomic data that currently are generated require flexible methods that can evolve with the data, so that they can help us answer important questions related to evolutionary processes in the wild. The field of genomic prediction has so far mainly been driven forward by animal and plant breeders, as well as by researchers in human genomics (Meuwissen et al., 2001; Wray et al., 2007). However, the current emergence of machine learning techniques relies on large data sets (Montesinos-López et al., 2021; Nazzicari and Biscarini, 2022; Gill et al., 2022), but sample sizes for wild populations are currently still too small to fully benefit from those latest advances. At the same time, many data sets are now growing to a critical volume that makes it more difficult to handle them with existing Bayesian methods, and computation times are becoming long (*e.g.*, Meher et al., 2022). We thus see BPCRR as a complementing alternative and a promising approach that fills an opening gap to handle current and future challenges of wild study systems.

## Acknowledgements

We are grateful to the many researchers, students and fieldworkers who helped collect the empirical data on house sparrows, laboratory technicians for assistance with laboratory analyses, and the local people in the study meta-population for their hospitality. Our study was supported by grants from the Norwegian Research Council (projects 274930 and 302619) and its Centre of Excellence funding scheme (project 223257). Genotyping on the custom house sparrow Affymetrix Axiom 200K SNP array was carried out at CIGENE, Norwegian University of Life Sciences, Norway. Computations were performed on resources provided by the NTNU IDUN/EPIC computing cluster (Själander et al., 2019).

## Appendix

## Appendix A: Two-step procedure in BayesR

For comparison, we ran a 10-fold cross-validation for each trait, and estimated posterior distributions for *V_A_* from the full data set using the BayesR v2 software package based on Moser et al. (2015). We used Dirichlet priors with 1, 1, 1, 5 and default values otherwise. All MCMC chains used to generate Figure S1 were run for 5000 iterations, a burn-in of 1000 and a saving frequency of 10. In all cases, computation time was between 1.5 and 3 hours on personal laptop with an Intel Core i7-1260P CPU. Note that these are relatively short chains compared to those generated by hibayes in the main text. Visual convergence checks indicate that the chains are stationary, but should be run 5-10 times longer if the aim was to derive reliable inference.

Here, the aim was mainly to illustrate the consequence of using a two-step procedure with BayesR (Figure S1). In a two-step approach, a linear mixed model (LMM) is first fitted with all the fixed and random effects except the genetic value. The estimated random individual-specific (ID) effect from this pre-fitted LMM is then used as the new response. However, while such a procedure leads to good prediction accuracy (Figure S1, top), it underestimates the actual variance in the genetic values, that is, *V_A_* (Figure S1, bottom a) - c)). The problem is particularly pronounced for body mass and wing length, where using the sum of the ID effect plus the mean of the residuals over all repeats within an individual (ID+res) recovers a correct *V_A_* estimate. Both body mass and wing length are plastic traits, thus the residuals absorb part of the within-individual variability of the genetic value. Tarsus length, on the other hand, is static skeletal trait and seems not (or much less) affected by the observed effect.

**Figure S1:**
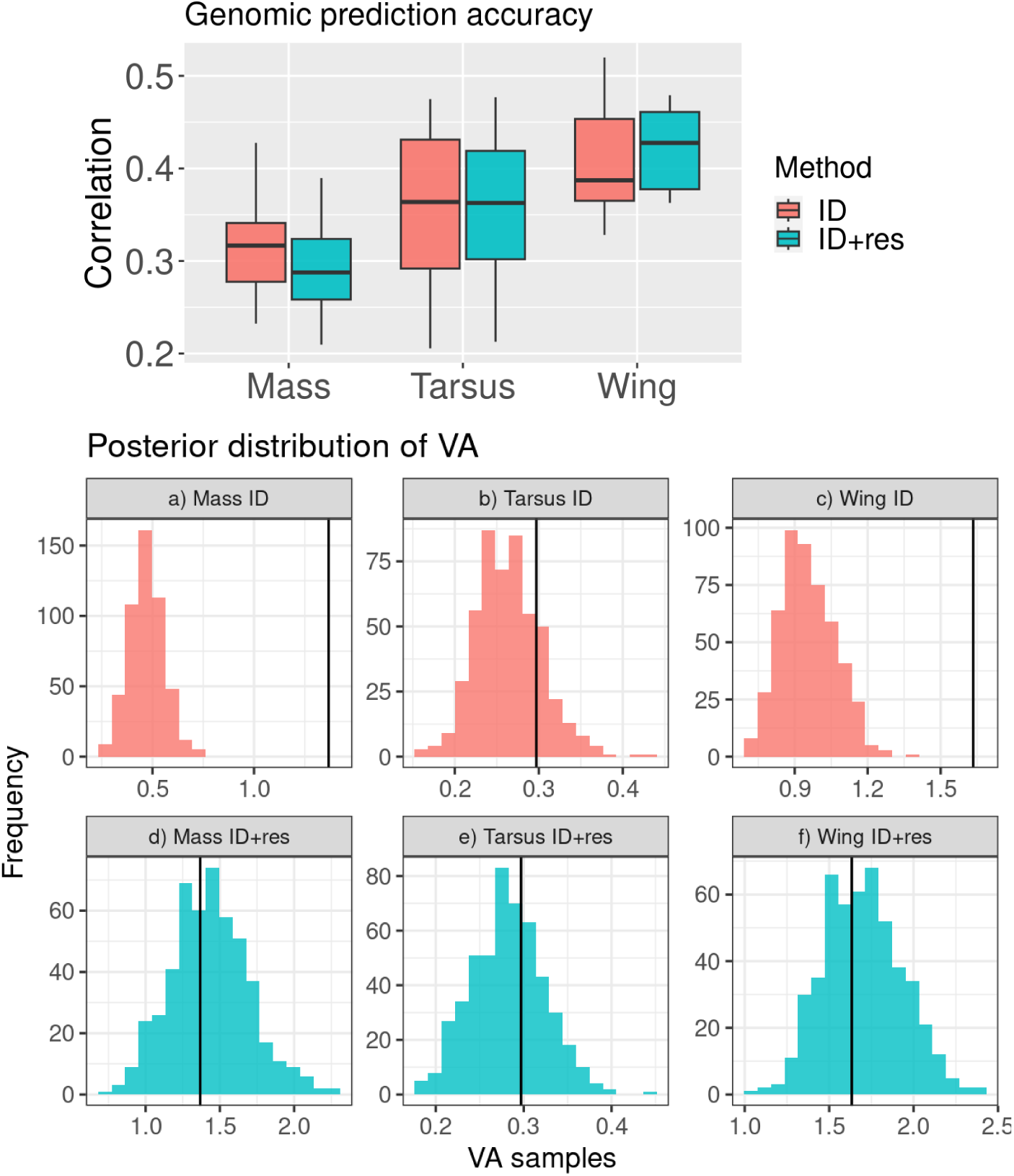
Comparison of the results for the two-step implementation of BayesR for the cases where the individual-specific ID effect is used as the new response in the second step (denoted as ID in the figure), and the case where we take the sum of the ID effect plus the mean of the residuals over all repeats within an individual (denoted as ID+res). The results from a 10-fold cross-validation illustrate that prediction accuracy tends to be higher when using ID as the new response (top), in particular for the two plastic traits mass and wing. However, *V_A_* then tends to be underestimated for those traits (bottom), while using ID+res, recovers correct *V_A_*. Vertical black lines indicate the estimated *V_A_* value from the genomic animal model.

**Figure S2:**
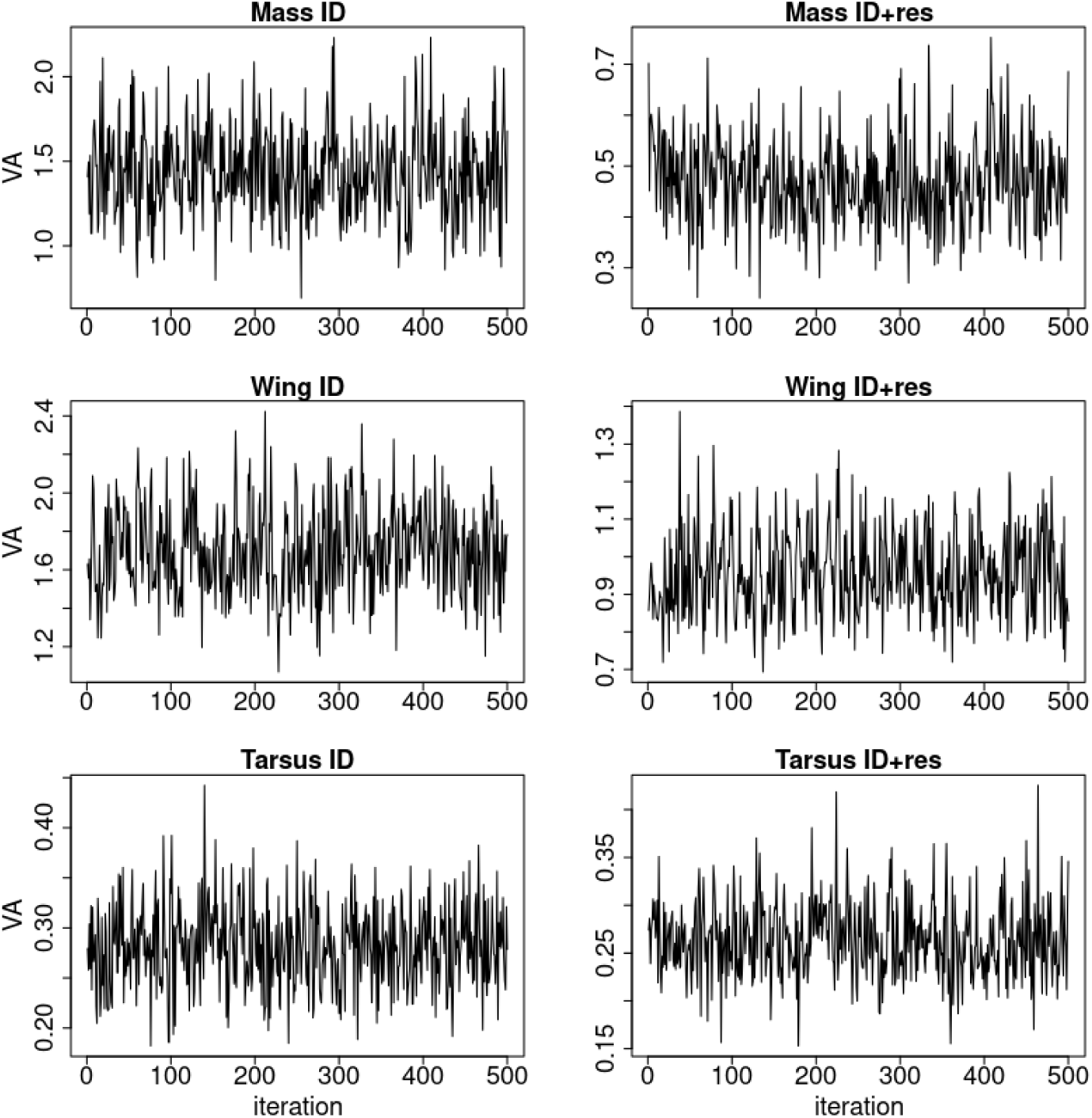
Trace plots for the six models analyzed with a two-step procedure, using the BayesR v2 software. Given that the thinning was 10, only every 10th MCMC iteration is shown.

## Appendix B: Proportionality of explained variances

If the infinitesimal model assumption holds, that is, for highly polygenic traits, we expect the variance explained by a PC to be linearly related to the variance it explains in the breeding value of the trait of interest. Another direct implication of the infinitesimal model assumption is that there is a linear correspondence between the proportion of variance explained by the first *k* PCs and the proportion of additive genetic variance explained in the trait of interest, which in turn corresponds to the relation between the estimated and true heritabilities 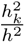 obtained from using the *k* first PCs. The approach for choosing the optimum number of PCs in fact relies on this assumption. Figure S3 indicates that the respective assumption is approximately fulfilled.

**Figure S3:**
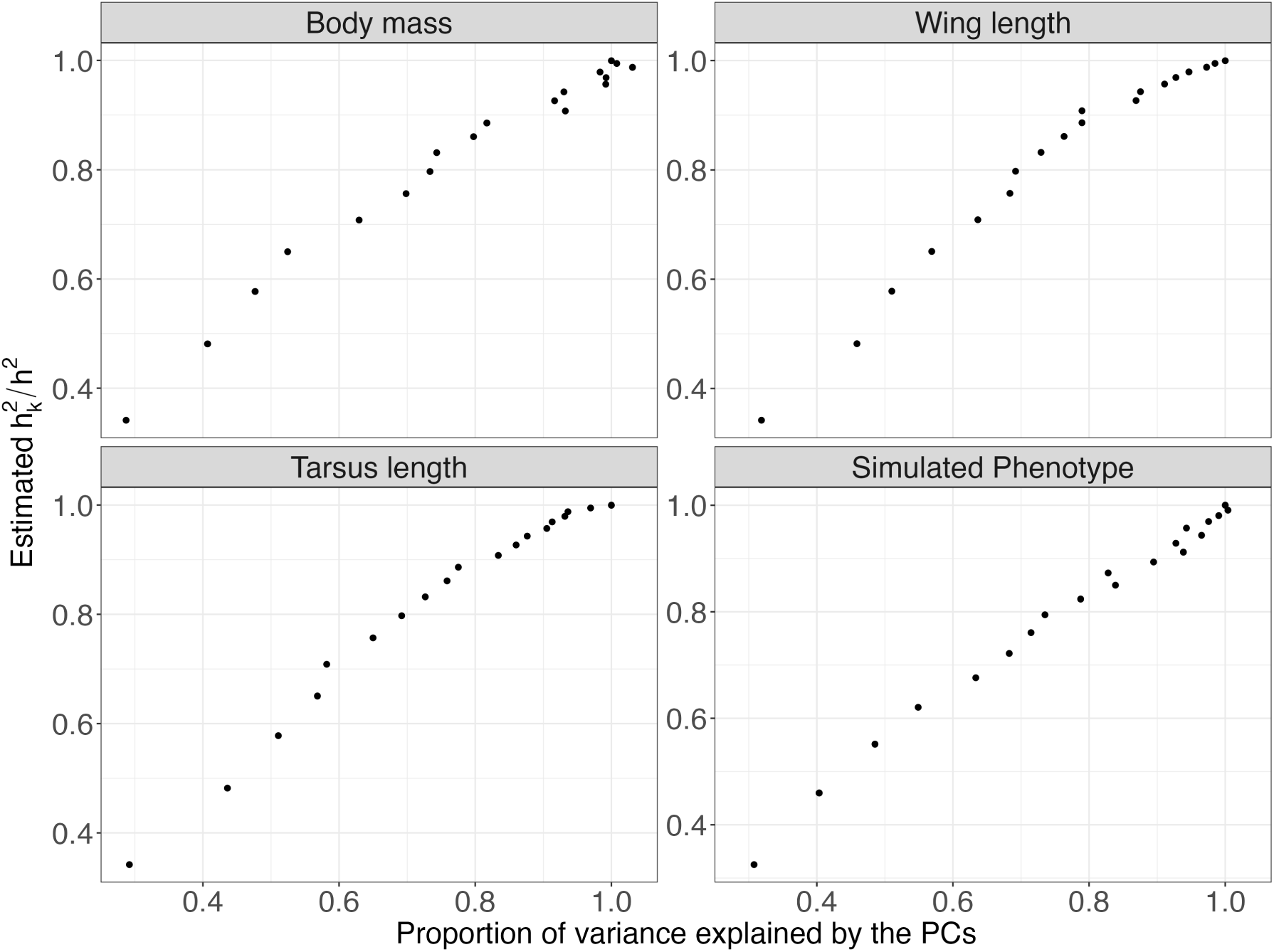
Relation between the proportion of estimated heritabilty 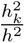 against the proportion of variance explained by the respective set of PCs. The results for the three traits (body mass, wing length and tarsus length) and the simulated phenotype indicate that the relationship is approximately linear, reflecting that the variance explained by a PC can be assumed approximately linearly related to the variance it explains in the breeding value of the trait of interest.

## Appendix C: Genomic prediction accuracy and its dependence on priors and scaling

In the main text (Figure 2), we have shown genomic prediction accuracies depending on the number of PCs for the case where the informative point priors from formula (4) were used. Importantly, the results look almost identical for the case where no prior knowledge on *σ_G_*^2^ was assumed, that is, when default priors 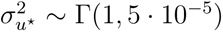 for the variance of the PC effects were used (Figure S4, green boxplots).

In addition, we have also compared different ways to scale the variances of the PCs. As described in the main text (Sections 2.2.2 to 2.2.4), we propose to scale the PC variances such that they remain proportional to their eigenvalues (Macciotta et al., 2010). A major benefit of not standardizing the variances to unity – which corresponds to the common ridge regression standardization for arbitrary predictors – is that even including large numbers of PCs then typically does not lead to significantly lower prediction accuracy than for the “best” *k* found by equation (6) in the main text, namely because PC-effects for PCs with less variance are automatically shrunken more, and over-fitting is omitted (Figure S4, green boxplots). In contrast, here we illustrate how standardizing the variances of all PCs to unity leads to a prediction accuracy decreases after the optimal balance is reached, as expected (Figure S4, blue boxplots).

**Figure S4:**
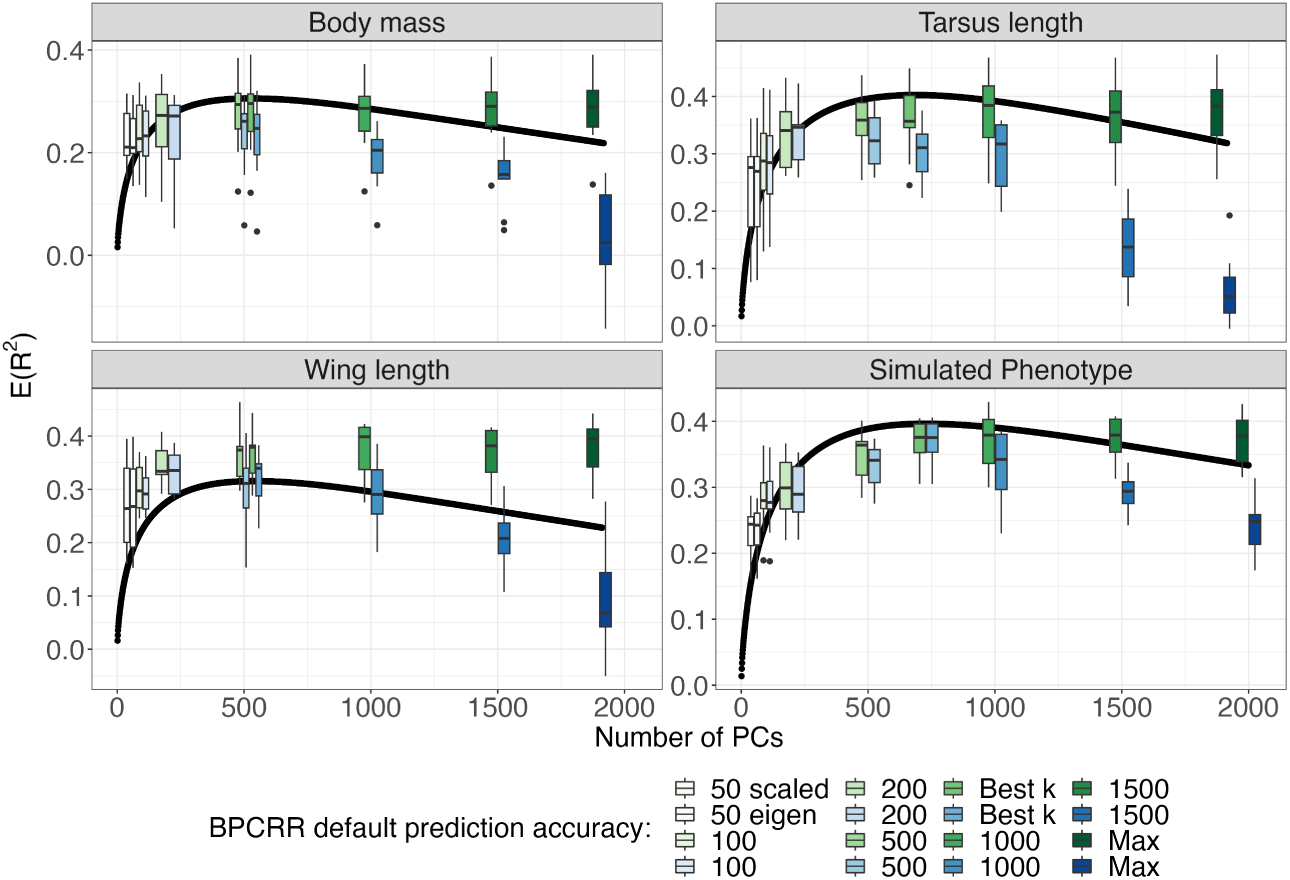
Expected prediction accuracy *E*(*R*^2^) using formulas (6) and (7), in dependence of the number of PCs (*k*) and for *h*^2^ and *N* values that correspond to the respective cases (black line). The actually observed prediction accuracies from a 10-fold cross-validation are added as boxplots for comparison (in green). The figure are enriched with results where all PCs are scaled to 1 (blue). The observed accuracies were obtained by using default priors in INLA.

## Appendix D: Convergence plots for hibayes

**Figure S5:**
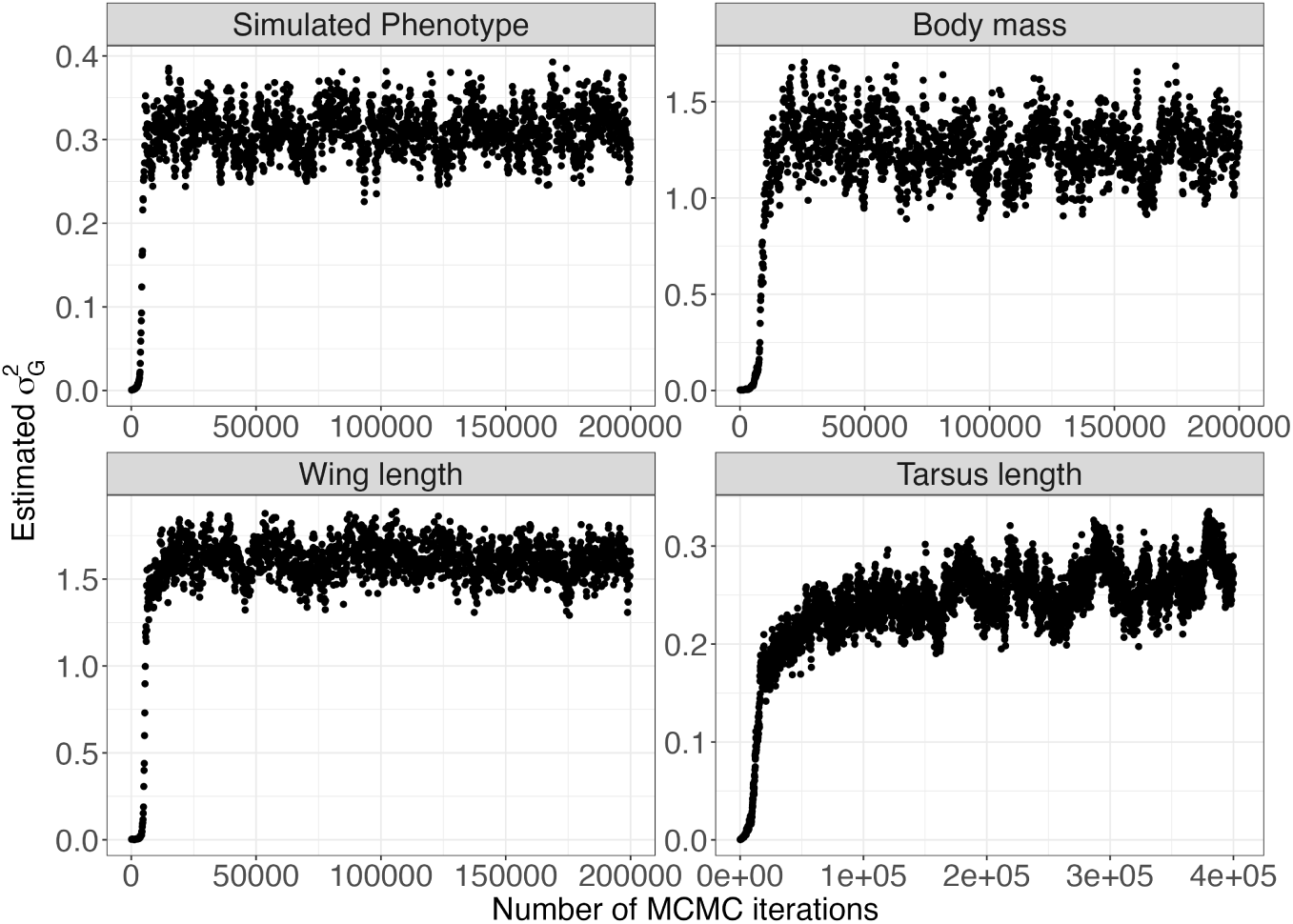
Convergence plots for simulated data and the traits for the longest chains that were run on the full dataset, with one data point for every hundred MCMC iterations. Note that tarsus length shows very slow convergence, and that even 400’000 iterations are a lower limit for this trait.

